# Phenotype loss is associated with widespread divergence of the gene regulatory landscape in evolution

**DOI:** 10.1101/238634

**Authors:** Juliana G. Roscito, Katrin Sameith, Genis Parra, Bjoern E. Langer, Andreas Petzold, Miguel Trefaut Rodrigues, Michael Hiller

## Abstract

Detecting the genomic changes underlying phenotypic changes between species is a main goal of evolutionary biology and genomics. Evolutionary theory predicts that changes in *cis*-regulatory elements are important for morphological changes. Here, we combine genome sequencing and functional genomics with genome-wide comparative analyses to investigate the fate of regulatory elements in lineages that lost morphological traits. We first show that limb loss in snakes is associated with widespread divergence of limb regulatory elements. We next show that eye degeneration in subterranean mammals is associated with widespread divergence of eye regulatory elements. In both cases, sequence divergence results in an extensive loss of relevant transcription factor binding sites. Importantly, diverged regulatory elements are associated with key genes required for normal limb patterning or normal eye development and function, suggesting that regulatory divergence contributed to the loss of these phenotypes. Together, our results provide the first evidence that genome-wide decay of the phenotype-specific *cis*-regulatory landscape is a hallmark of lost morphological traits.

## Introduction

Phenotypic diversity is most easily observable as differences in morphology. Understanding how morphological differences evolved is a central question in many areas of biology. Morphology is established during development and requires the patterning of the developing embryo to assign specific fates to cells. Patterning processes are controlled by genes that are often involved in the development of many different structures. Consequently, expression of these highly pleiotropic developmental genes must be tightly regulated in a spatio-temporal manner. Key to transcriptional regulation of pleiotropic developmental genes are *cis*-regulatory elements that can be located far upstream or downstream of the promoter and often control gene expression in specific tissues at specific time points. Thus, understanding how developmental genes and their *cis*-regulatory elements evolve is crucial to understanding how morphology evolves.

Differences in pleiotropy between developmental genes and *cis*-regulatory elements impact which mutations are permissible in evolution. Whereas mutations in the coding regions of a pleiotropic gene may affect gene function in many tissues, which is often deleterious, mutations in modular *cis*-regulatory elements likely affect gene expression only at a specific time and tissue. Accordingly, it was proposed that morphology largely evolves by changes in the spatio-temporal expression of developmental genes, which in turn evolves by changes in the underlying *cis*-regulatory elements^1,2^. This hypothesis is supported by a growing body of evidence^3–21^.

The loss of a complex phenotype is one extreme case of morphological evolution. Upon phenotype loss, we expect a different evolutionary trajectory for the genetic information underlying this phenotype. On the one hand, the integrity of developmental genes should be maintained over time due to selection on those gene functions that are not related to the lost phenotype. On the other hand, modular *cis*-regulatory elements associated specifically with this phenotype may directly contribute to its loss and are generally expected to evolve neutrally afterwards. This should result in sequence divergence and thus decay of regulatory activity over time. These different trajectories are illustrated by the loss of pelvic spines in freshwater stickleback fish, caused by the loss of a pelvic-specific enhancer for the pleiotropic *Pitx1* gene, while *Pitx1* remained intact^10^. Similarly, mutations in the limb-specific ZRS enhancer for sonic hedgehog (Shh) in the snake lineage led to altered *Shh* expression in the limbs, while the pleiotropic *Shh* gene remained intact^17,18^. However, recent studies^18,22,23^ found that numerous other limb enhancers are nevertheless still conserved in snakes, despite limb reduction in this lineage dating back to more than 100 Mya^24^. Thus, it remains an open question whether phenotype loss is generally associated with divergence of the *cis*-regulatory landscape on a genome-wide scale.

Here, we combine genome sequencing with functional and comparative genomics to systematically investigate the fate of *cis*-regulatory elements in lineages that lost complex phenotypes: loss of limbs in snakes and degeneration of eyes in subterranean mammals. Our analyses provide the first genome-wide evidence that divergence of the phenotype-specific *cis*-regulatory landscape is a hallmark of lost morphological traits. Furthermore, we present the first comprehensive picture of the non-coding genomic changes that likely contributed to limb loss and eye degeneration. More generally, our study provides a widely applicable comparative and functional genomics strategy to detect *cis*-regulatory element candidates that may underlie morphological differences between species.

## Results

### Genome sequencing and genome comparison between snakes and limbed vertebrates

We first investigated the fate of the limb-related *cis*-regulatory landscape in snakes. To detect sequence divergence that is specific to snakes, it is necessary to compare the genomes of snakes to the genomes of several fully limbed reptiles and other vertebrates. Given the sparsity of genomes of reptiles with well-developed limbs, we sequenced and assembled the genome of the fully limbed tegu lizard *Salvator merianae*, representing the first sequenced species of the teiid lineage. We used a combination of Illumina MiSeq and HiSeq sequencing (Supplementary Table 1), followed by iterative read error-correction with SGA-ICE^25^, and genome assembly with ALLPATHS-LG^26^. The resulting 2 Gb genome assembly has a contig N50 value of 176 Kb and a scaffold N50 value of 28 Mb, with the longest scaffold spanning 99 Mb. Compared with other sequenced reptiles, the tegu genome shows the largest contig N50 value and the second largest scaffold N50 value (Figure 1A). Our tegu assembly contains 196 of the 197 vertebrate non-exonic ultraconserved elements^27^, showing an assembly completeness comparable to or better than that of all other sequenced reptiles (Supplementary Table 2). Combining transcriptomics and homology-based gene prediction approaches, we annotated a total of 23,487 genes, 16,284 of which have a human ortholog (Methods). Using the tegu genome as reference, we created a multiple genome alignment including two well-assembled snakes (boa and python), three other limbed reptiles (green anole lizard, dragon lizard, and gecko), three birds, alligator, three turtles, 14 mammals, frog and coelacanth (Figure 1B; Supplementary Table 3; Methods). This alignment of 29 genomes was used as the basis for all further analyses.

**Figure 1:**
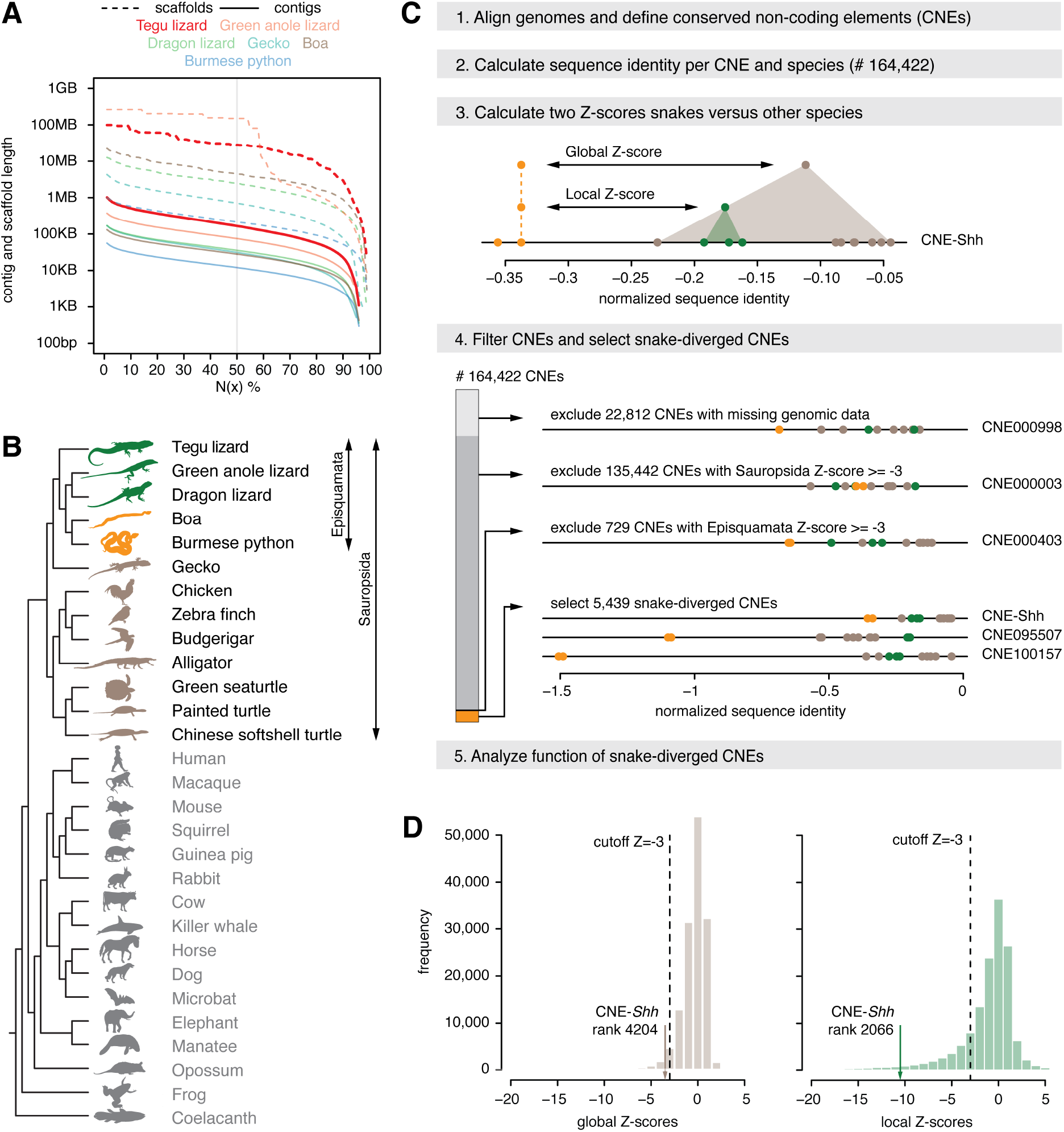
Comparative framework to detect sequence divergence of regulatory elements in snakes on a genome-wide scale. (A) Comparison of the tegu genome to genomes of other sequenced reptiles. N(x)% graph showing the contig and scaffold size (y-axis), where x% of the genome consists of contigs and scaffolds of at least that size. (B) Phylogenetic tree of the limbless and limbed species included in our multiple genome alignment. Snakes are in orange, remaining Episquamata species in green, remaining Sauropsida species in brown, and outgroup species in grey. (C) Overview of the computational steps to identify CNEs that are specifically diverged in snakes. (D) Histograms depicting global and local Z-score distributions of 141,610 CNEs (22,812 CNEs with missing genomic sequence are excluded).

### Thousands of conserved non-coding elements are specifically diverged in snakes

To study the evolution of *cis*-regulatory elements across species on a genome-wide scale, we focused on conserved non-coding elements (CNEs), because evolutionary sequence conservation implies purifying selection and thus function, and because CNEs often overlap *cis*-regulatory elements^28–31^. Using our genome alignment, we identified a total of 164,422 non-coding elements (covering 1% of the tegu genome) that are conserved among many but not necessarily all Amniota species (workflow in Figure 1C). For each CNE, we computed a per-species sequence divergence value by determining the percent of bases that are identical between the species’ CNE sequence and the reconstructed sequence of the amniote ancestor. By searching for CNEs that exhibit a substantially lower sequence identity in both boa and python snakes, we expect to identify *cis*-regulatory elements that control gene expression in the developing limbs of limbed species and that may no longer function as limb regulatory elements in limbless snakes. To identify such snake-diverged CNEs, we first computed a global Z-score, which measures how many standard deviations the sequence identity in snakes is below the average identity in all other limbed species. To assure that divergence is specific to snakes and to exclude relaxed selection in the entire reptile clade, we additionally computed a local Z-score by comparing the snakes and the three lizards alone. Requiring a Z-score cutoff of −3 for both comparisons, we identified 5,439 CNEs that are highly and specifically diverged in snakes (Supplementary Table 4). Importantly, this set includes the well-studied ZRS limb enhancer that likely underlies *Shh* mis-expression in snakes^17,18^ (Supplementary Figure 1). However, sorted by the global Z-score, the *Shh* enhancer ranks at position 4,204 (Figure 1D), showing that numerous other CNEs, including CNEs near key limb developmental genes (see below; Supplementary Figure 2), have a more striking snake-specific divergence pattern.

### Snake-diverged CNEs are significantly associated with limb-related genes

If snake-diverged CNEs function as limb regulatory elements, we expect that they should preferentially be located near genes with a known role in limb development. To test this, we first determined potential target genes for each CNE. Using an approach similar to GREAT^32^, we associated each CNE to the closest neighboring genes that are located up to 300 Kb up- or downstream of the CNE (Supplementary Figure 3). Next, we assessed whether and to what extent snake-diverged CNEs are preferentially associated with limb-related genes, compared with the remaining CNEs which are not diverged in snakes.

The analysis shows that snake-diverged CNEs are significantly enriched near genes that are involved in limb development and linked to congenital limb malformations^33^ (Figure 2A, top panel; Supplementary Table 5). This includes *Tbx4, En1, Gli3, Grem1, Hand2*, and other genes that play important roles in limb patterning and outgrowth (see below). Furthermore, we found a significant association with genes whose knockouts in mouse result in limb phenotypes^34^ (Figure 2A, bottom panel; Supplementary Table 6). For example, 54 snake-diverged CNEs are associated with 11 genes whose knockout results in an abnormal apical ectodermal ridge, an essential signaling center required for limb outgrowth. Interestingly, the snake-diverged CNEs that are associated with limb-related genes are predominantly located outside of promoter-proximal regions (Figure 2A, right panel), indicating divergence of distal *cis*-regulatory elements.

**Figure 2:**
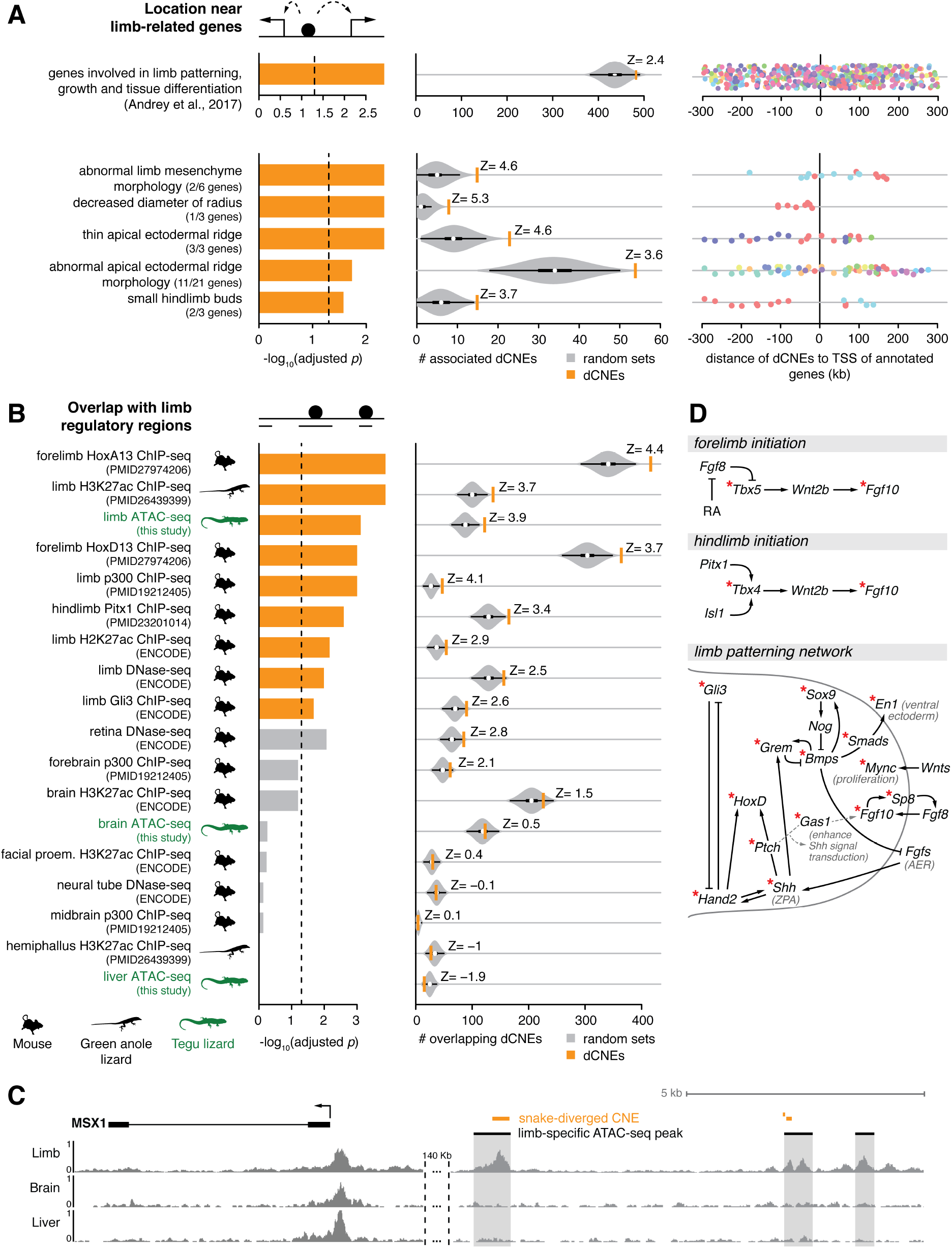
Snake-diverged CNEs are associated with limb-related genes and significantly overlap limb regulatory elements. (A) Snake-diverged CNEs are significantly associated with genes annotated with limb-related functions. Left: Bars depict adjusted p-values derived by a one-sided Fisher’s exact test. Middle: Observed (orange vertical bar) and expected number (grey violin plots, based on 10,000 random subsets sampled from all CNEs) of snake-diverged CNEs associated with genes in each set (the thick box inside the violin plot indicates the first quartile, the median and the third quartile). The Z-score measuring the number of standard deviations that the observed number is above the random expectation is indicated. Right panel: Many snake-diverged CNEs (dots) are far away from the transcription start site of genes in these sets. CNEs associated with the same gene have the same color. (B) Snake-diverged CNEs significantly overlap regulatory elements active in embryonic limb tissue of tegu lizard, green anole lizard, and mouse. Orange bars correspond to limb regulatory datasets. ATAC-seq datasets generated in this study are highlighted in green. Remaining visual representation as in (A). (C) Genome browser screenshot shows that snake-diverged CNEs (orange) overlap tegu limb-specific ATAC-seq peaks. ATAC-seq signal tracks of limb, brain and liver are shown. (D) Snake-diverged CNEs are broadly distributed in the limb patterning network. An asterisk marks the genes which are associated with at least one snake-diverged CNE that overlaps a limb regulatory element.

To confirm that these enrichments are specific to snake-diverged CNEs, we obtained CNEs diverged in the sister lineage comprising the limbed anole and dragon lizards, repeated the statistical tests and observed no significant association with limb-related genes (Supplementary Table 6). Taken together, these results show that the limb-related enrichments are specific to CNEs diverged in snakes, indicating that mutations in these CNEs could affect expression of important limb developmental genes.

### Snake-diverged CNEs significantly overlap limb regulatory elements

To directly test if snake-diverged CNEs overlap *cis*-regulatory elements that are active during normal limb development in species with fully developed limbs, we obtained embryonic limb tissue of the tegu lizard and used ATAC-seq^35^ to identify regions of accessible chromatin, which often correspond to regions with gene regulatory activity^36,37^. To assess tissue-specificity of limb regulatory elements, we additionally determined accessible chromatin in tegu lizard brain, heart and liver tissues, the lateral flank tissue between limb buds, and tissue of the remaining embryo. We identified 5,635 limb-specific ATAC-seq peaks in the tegu lizard (Figure 2C). These limb-specific peaks are significantly associated with genes involved in limb development (GREAT analysis^32^, Supplementary Figure 4) and overlap many known limb enhancers for *Fgf10*^38^, *Grem1*^39,40^, *Prrx1*^41^, *Bmp4*^42^, *Bmp2*^43^, *HoxA*^44^, *Twist1*^45^, *Shh*^46^, *Sox8*^47^, and other genes^48^ (Supplementary Figure 5, Supplementary Table 7). Similarly, the 5,417 brain-specific peaks and 6,112 liver-specific peaks are significantly associated with genes having brain and liver functions, respectively (Supplementary Figure 4). These results validate the specificity of the ATAC-seq signal.

We found a highly significant overlap between snake-diverged CNEs and the tegu limb-specific ATAC-seq peaks (green labels in Figure 2B), in comparison with the remaining non-diverged CNEs (Supplementary Table 8). To further corroborate this observation, we compiled a rich list of publicly available limb regulatory datasets from two other limbed species (mouse and green anole lizard), mapped the data to the tegu genome, and statistically tested the significance of the overlap (Supplementary Table 8). We found a significant overlap between snake-diverged CNEs and several limb datasets, such as limb-specific H3K27ac enhancer marks^22,49^ and accessible chromatin obtained with DNase-seq^49^. Snake-diverged CNEs also significantly overlap limb-specific binding sites of the transcriptional co-activator p300^50^, and binding sites of the limb-related transcription factors HOXA13^44^, HOXD13^51^, PITX1^52^ and GLI3^40^ (Figure 2B). In total, we found 933 snake-diverged CNEs that overlap any of the tested limb regulatory datasets (Supplementary Table 4).

Notably, snake-diverged CNEs overlap many experimentally characterized limb enhancers^48^ (Supplementary Table 4), including enhancers regulating *HoxA* genes and the *HoxD* enhancers CsB and Island I^44,53,54^. In addition to the well-known *Shh* ZRS limb enhancer whose divergence likely underlies mis-expression of *Shh* in snake limb buds^17,18,55^, snake-diverged CNEs overlap a VISTA limb enhancer near *Gli3*, a transcription factor that represses *Shh* in the limb bud^56,57^. We also detected snake-specific sequence divergence in a VISTA limb enhancer near *Gas1*, a gene encoding a membrane protein that enhances *Shh* signaling and that is required for the normal regulatory loop between *FGF10* in the mesenchyme and *FGF8* in the apical ectodermal ridge^58,59^. This suggests a broader divergence in the *Shh* signaling pathway, including components which affect both expression and read-out of this important signaling morphogen.

To confirm that these enrichments are specific to limb regulatory elements, we used our ATAC-seq and other publicly available regulatory data from non-limb tissue and found no significant overlap with snake-diverged CNEs (Figure 2B; Supplementary Table 8), with the exception of eye regulatory data (see Discussion). In addition, we repeated the analysis using the set of CNEs diverged in the green anole and dragon lizard, and found no significant overlap with limb datasets (Supplementary Table 8), showing that the association with limb regulatory elements is specific to CNEs diverged in snakes. These results provide strong evidence that many snake-diverged CNEs overlap regulatory elements that control the expression of important limb-related genes in other limbed animals.

### Snake-specific CNE divergence is widespread across the limb patterning network

As snake-diverged CNEs are preferentially associated with limb-related genes and overlap limb regulatory elements, we assessed whether snake-specific regulatory divergence is limited to specific signaling pathways, or whether divergence is widespread across the limb developmental patterning network. To this end, we compiled a limb regulatory network based on^60–63^ and highlighted the genes that are associated with the 933 snake-diverged CNEs that overlap limb regulatory elements (Supplementary Table 4). This analysis shows a global pattern of divergence affecting all the major limb signaling pathways (Figure 2D). Notably, snake-diverged CNEs are associated with key genes controlling fore- and hindlimb initiation, anterior-posterior and dorsal-ventral patterning, and limb outgrowth, such as *Tbx4/5, Fgf10, Hand2, HoxD, Grem1, Shh, Gli3, Gas1, Sox9, Bmp4, En1*, and others. In addition to *Shh*, many of these genes cause severe limb truncations and other limb malformations when knocked-out or when their spatio-temporal expression is perturbed^17,18,64-73^. Furthermore, genes such as *Shh, Gli3*, and *Hand2* were shown to be mis-expressed in the limb buds of developing snakes^18,55^. Together, this suggests that snake-specific divergence in many limb regulatory elements may have contributed to the loss of limbs in the evolutionary history of snakes.

### Eye degeneration in subterranean mammals allows to assess whether regulatory divergence is a general evolutionary principle

The compelling evidence for genome-wide divergence of the limb regulatory landscape in snakes raises the question if this phenomenon is specific to limb loss or rather a general evolutionary principle. To address this question, we analyzed the fate of eye regulatory elements in four subterranean mammals in which eye degeneration evolved independently: the blind mole rat, naked mole rat, star-nosed mole, and cape golden mole. These species exhibit reduced eye sizes, disorganized lenses, thinner retinas, and loss of optic nerve connections^74–78^. A recent study has shown that a number of eye enhancers are evolving at accelerated rates in these four vision-impaired species^79^; however, the divergence of eye regulatory elements was not assessed on a genome-wide basis.

To investigate if eye degeneration is associated with genome-wide divergence of the eye regulatory landscape in the vision-impaired subterranean mammals, we conducted a comparative analysis contrasting these species with mammals that do not have a degenerated visual system. Similar to our analysis of limb loss in snakes, we created a multiple genome alignment, using mouse as the reference and including the genomes of the aforementioned four subterranean mammals, 16 other mammalian species, the green anole lizard, frog and chicken (Figure 3A; Supplementary Table 9). With this alignment of 24 genomes, we identified 491,576 CNEs (covering 3.3% of the mouse genome), and measured sequence identity per species with respect to the reconstructed placental mammal ancestor. As the degeneration of the visual system evolved independently in the four subterranean lineages, we identified CNEs diverged in these mammals with the previously developed “Forward Genomics” branch method that considers phylogenetic relatedness between species^80^. Of all 491,576 CNEs, this method identified 9,364 CNEs with significantly higher sequence divergence in the subterranean mammals compared to the other mammals (adjusted *p*-value cutoff 0.005, Supplementary Figure 6; Supplementary Table 10).

**Figure 3:**
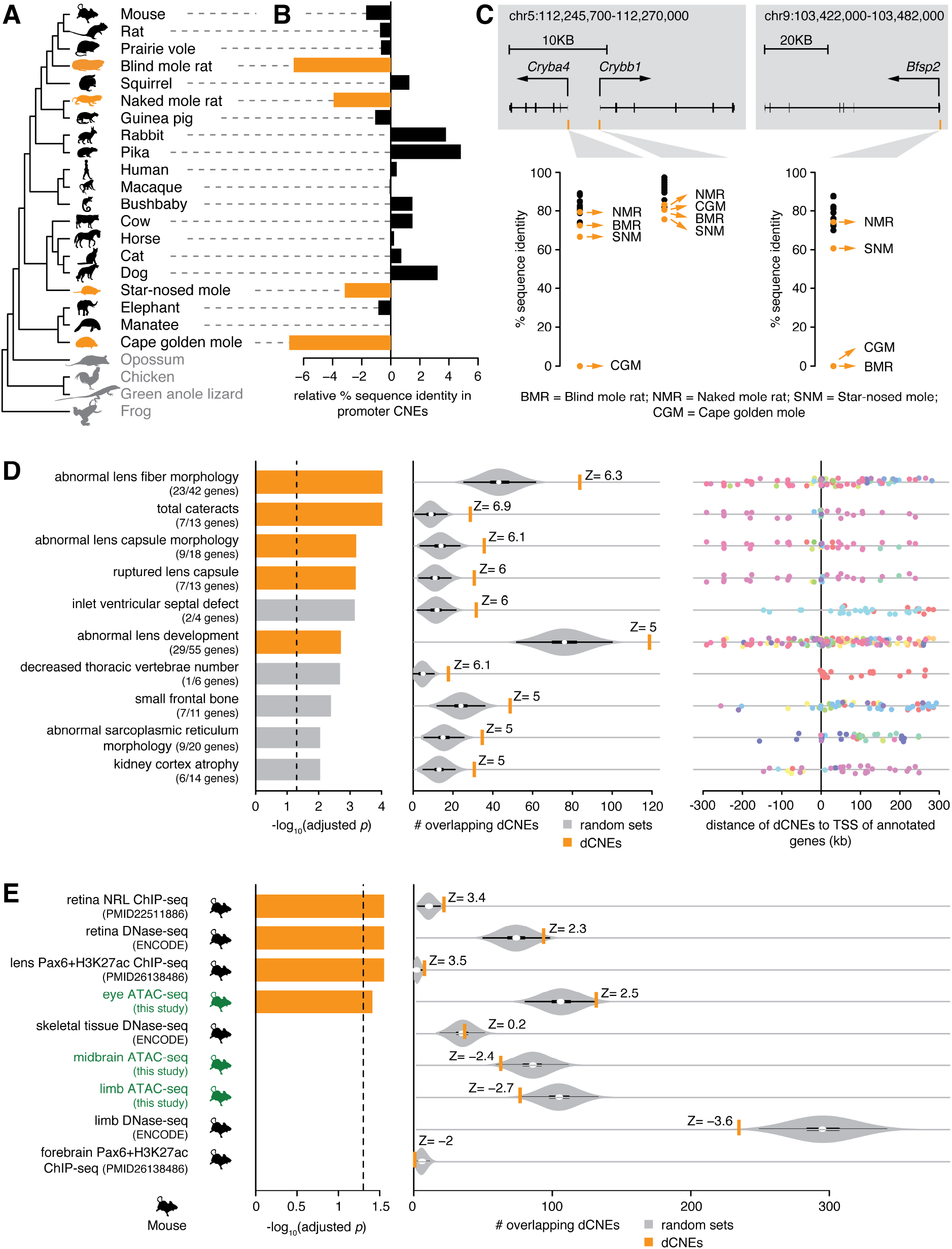
CNEs specifically diverged in vision-impaired subterranean mammals are associated with eye-related genes and significantly overlap eye regulatory elements. (A) Phylogenetic tree of species in our multiple genome alignment. Subterranean mammals are in orange, mammals without degenerated eyes are in black, and outgroup species are in grey. (B) Difference in the mean percent sequence identity between 52 CNEs overlapping the promoter regions of 64 eye-specific genes, which are diverged or lost in subterranean mammals, and 18,033 CNEs overlapping promoters of other genes. Compared to other mammals, the subterranean mammals have substantially higher divergence in the CNEs overlapping the 64 eye-specific gene promoters. Promoters are defined as ±1.5 Kb around the transcription start site. (C) Three examples of diverged CNEs located in the promoter of diverged lens-specific genes. (D) Diverged CNEs are enriched near genes that lead to lens defects in a mouse knockout (top 10 enrichments are shown here; Supplementary Table 12 lists all enrichments). Lens-related knockout phenotypes are shown in orange. Remaining visual representation as in Figure 2A.(E) Diverged CNEs significantly overlap regulatory elements active in whole eye, retina and lens. Orange bars indicate eye regulatory datasets. ATAC-seq datasets generated in this study are highlighted in green. Remaining visual representation as in Figure 2B.

### CNEs diverged in subterranean mammals are significantly associated with promoters of diverged eye-related genes

To explore if these 9,364 CNEs are associated with eye-related genes, we first focused on 64 eye-specific genes that are diverged or lost in at least one of the four subterranean mammals, according to previous studies^80–83^ (Supplementary Table 11). Since relaxed selection on these genes should result in promoter divergence, these 64 promoter regions should be enriched in CNEs diverged in subterranean mammals. Indeed, 23% (12/52) of the CNEs located in the 64 promoters were identified in our genome-wide screen for divergence in subterranean mammals, which is a significant enrichment compared to 2% (357/18,033) of the CNEs located in the promoters of all other genes (Fisher’s exact test: *p* = 4.3e^−10^). Consistently, the average sequence divergence of the 52 CNEs located in these 64 promoter regions is substantially higher in the subterranean mammals (Figure 3B). Examples illustrating promoter divergence in the lens-specific genes *Cryba4, Crybb1*, and *Bfsp2* are shown in Figure 3C. Overall, CNEs located in the promoter regions of diverged eye-related genes show the expected divergence pattern in subterranean mammals.

### CNEs diverged in subterranean mammals are significantly associated with eye-related genes

If divergence of the *cis*-regulatory landscape is a hallmark of lost morphological traits, we expect the CNEs diverged in subterranean mammals to be preferentially located near genes with a known role in eye development and function. Indeed, we found that diverged CNEs are significantly associated with genes whose knockout in mouse results in lens and cornea defects (Figure 3D; Supplementary Table 12). Importantly, excluding the aforementioned 64 diverged genes results in virtually the same enrichments (Supplementary Table 13). This shows that diverged CNEs are not only associated with these diverged genes, but also with many other genes encoding morphogens or transcription factors that are involved in optic cup, lens and retina development, for example *Atf4, Fgf9, Foxe3, Lhx2, Pax6, Prox1, Rax, Sox1*, and *Vsx2*. Furthermore, most of these CNEs are far away from the transcription start site (Figure 3D, right panel), indicating widespread divergence of distal *cis*-regulatory elements. We repeated this analysis for CNEs diverged in four mammals that do not have degenerated eyes (human, rat, cow, elephant), and found no significant association with eye-related genes (Supplementary Table 12). Taken together, these results show that the eye-related enrichments are specific to CNEs diverged in subterranean mammals.

### CNEs diverged in subterranean mammals significantly overlap eye regulatory elements

To directly test if CNEs diverged in subterranean mammals correspond to *cis*-regulatory elements active during normal eye development, we performed ATAC-seq^35^ on mouse whole eye at embryonic day E11.5, and lens and retina tissues at E14.5 (Supplementary Table 1). To assess tissue-specificity of eye regulatory elements, we also determined open chromatin in mouse limb and midbrain tissues at embryonic day E11.5. This analysis resulted in 8,020 eye-specific, 8,401 lens-specific, and 8,900 retina-specific mouse ATAC-seq peaks. These peaks are highly enriched near genes with annotated eye-, lens- and retina-related functions (Supplementary Figure 7). Furthermore, these peaks overlap characterized eye enhancers that regulate *Pax6*^84,85^ and other genes^48^ (Supplementary Figure 8, Supplementary Table 14). Likewise, 7,953 limb- and 5,779 midbrain-specific mouse ATAC-seq peaks are enriched near genes with limb and brain functions, respectively, validating the specificity of the ATAC-seq signal.

We found that CNEs diverged in subterranean mammals overlap tissue-specific ATAC-seq peaks detected in the developing mouse eye, lens, and retina significantly more often than expected from non-diverged CNEs (green labels in Figure 3E; Supplementary Table 15). In addition to these developmental datasets, we compiled a list of publicly available regulatory datasets from different eye tissues (Supplementary Table 15) and found significant overlap between diverged CNEs and several datasets, such as retina-specific DNase-seq peaks^49^, retina-specific binding sites of the transcription factor NRL^86^, and lens-specific enhancers bound by PAX6^87^. CNEs diverged in subterranean mammals also overlap VISTA eye enhancers (Supplementary Table 10), including an enhancer near *BMPER*, a gene that results in reduced eye size in a mouse knockout. In total, we found 575 diverged CNEs that overlap any of the tested eye regulatory datasets (Supplementary Table 10). In contrast, we found no significant overlap with our limb- and midbrain-specific ATAC-seq peaks, or other publicly available regulatory datasets from non-eye tissues (Figure 3E, Supplementary Table 15). Additionally, CNEs diverged in human, rat, cow and elephant showed no significant overlap with eye regulatory datasets (Supplementary Table 15).

In summary, we show that hundreds of CNEs diverged in subterranean mammals are located near genes with a role in eye development and function, and overlap regulatory elements that control gene expression during normal eye development and in adult eyes. Together with our equivalent results for snake-diverged CNEs, this corroborates that widespread divergence of the phenotype-specific *cis*-regulatory landscape is a hallmark of lost morphological traits.

### CNE divergence results in loss of transcription factor binding sites

If the sequence divergence that we consistently detected in snake-diverged CNEs and in CNEs diverged in subterranean mammals is a signature of functional decay, we expect that these diverged CNEs also lost binding sites of transcription factors (TFs) that have a role in limb and eye tissues. To test this, we first estimated the binding affinity of a TF to the reconstructed ancestral sequence of each CNE by computing a score based on the TF binding preference (motif). This motif score is calculated from the summed contribution of weak and strong binding sites (Methods) and makes no assumption about motif position, thus it is robust to binding site turnover. We validated this approach by obtaining available ChIP-seq experiments and showing that the genomic regions bound by a particular TF get high motif scores for this TF (Supplementary Figure 9). Then, we selected TF-CNE combinations where a TF motif is present in the reconstructed ancestral sequence, and determined the score of that particular motif in the corresponding CNE sequences of all descendant species. High motif scores imply motif conservation in the descendant species, while low scores imply motif loss. Finally, for each species, we compared the motif score distribution between diverged CNEs (foreground) and the remaining CNEs (background), considering a total of 55/58 motifs of TFs having a role in limb/eye development.

We first applied this strategy to compare the 933 snake-diverged CNEs that overlap limb regulatory data (Supplementary Table 4) with all remaining CNEs. For all limbed species, we found that the median motif score of these snake-diverged CNEs is significantly higher (adjusted *p* ≤ 7.0e^−16^, one-sided Wilcoxon rank sum test; Figure 4A). This shows that limb TF motifs are significantly conserved in limbed species, which is consistent with selection acting to preserve limb regulatory function. In sharp contrast, there is a clear absence of conservation of limb TF motifs in both snakes (adjusted *p* = 1), showing that snake-specific sequence divergence results in a large-scale loss of TF binding sites (Figure 4A and B).

**Figure 4:**
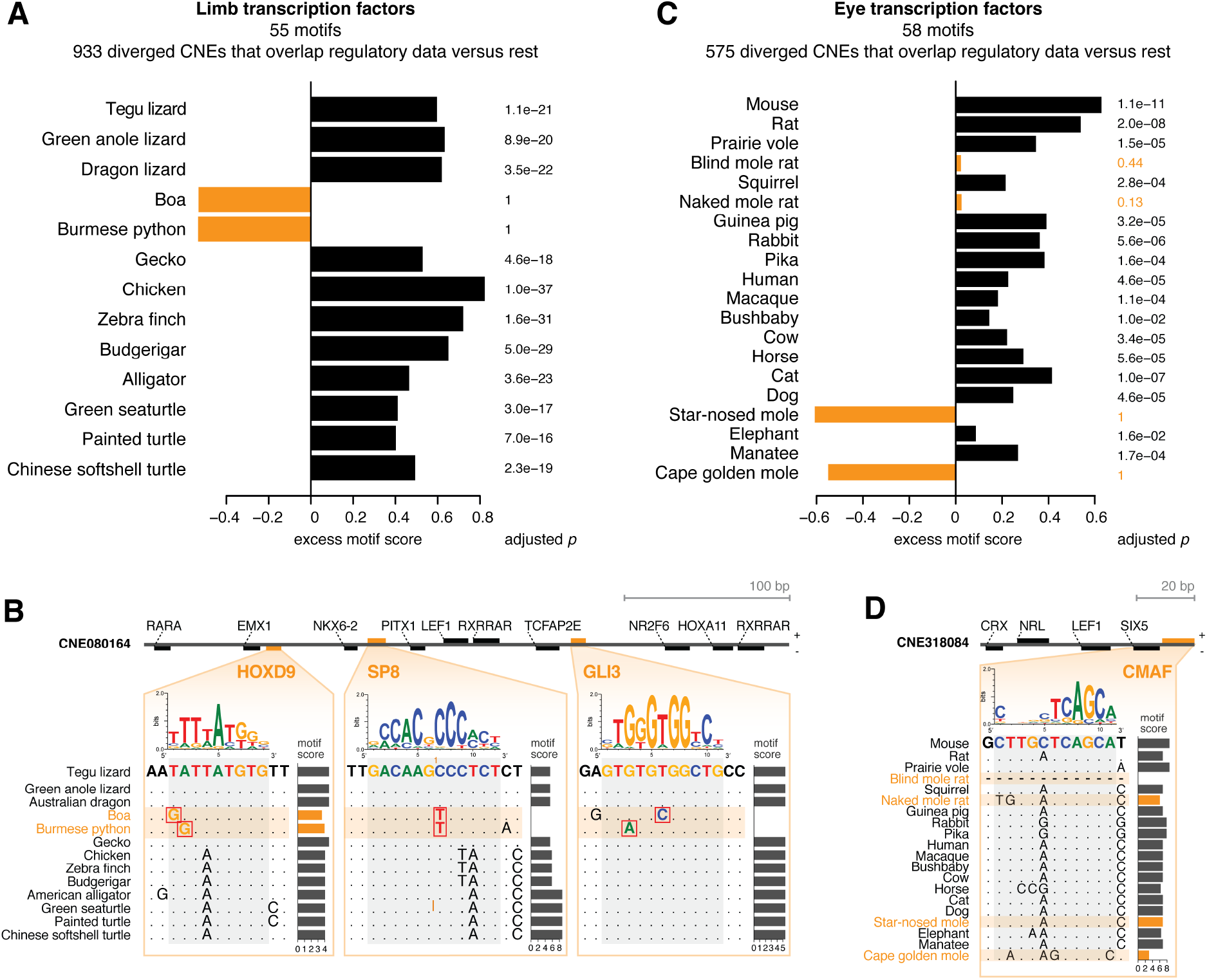
Sequence divergence results in widespread loss of binding sites of limb and eye-related transcription factors (TFs) in snakes and subterranean mammals. (A) Difference in the median motif scores of limb-related TFs between 933 snake-diverged CNEs that overlap limb regulatory data and all other CNEs (excess score, x-axis). A positive excess score as observed for limbed species reflects a preference to preserve motifs of limb-related TFs. P-values comparing the distribution of motif scores of diverged and all other CNEs were computed with a one-sided Wilcoxon rank sum test and corrected for multiple testing. (B) Mutations in snakes in a CNE overlapping a limb-specific ATAC-seq peak weaken or destroy binding sites for key limb transcription factors. The binding preference of the TFs is visualized as a sequence logo, motif scores were computed with SWAN^135^. (C) As panel A, but using motifs of eye-related TFs to compare CNEs that are diverged in subterranean mammals and overlap eye regulatory data with all other CNEs. (D) Mutations in two subterranean mammals in a CNE overlapping a lens-specific ATAC-seq peak weaken a binding site for c-Maf, a crystallin-inducing TF that is required for lens fiber cell differentiation^136,137^.

Next, we applied the same analysis to the 575 CNEs that are diverged in subterranean mammals and overlap eye regulatory data (Supplementary Table 10). Similarly, we found that in all species without degenerated eyes, eye TF motifs are significantly conserved (adjusted *p* < 1.6e^−2^; Figure 4C and D), in contrast to all four subterranean mammals (adjusted *p* ≥ 0.13). Altogether, this analysis implies that CNE divergence results in a large-scale divergence of relevant TF binding sites, supporting the observation that phenotype loss is associated with the genome-wide decay of the phenotype-specific *cis*-regulatory landscape.

## Discussion

Our combined comparative and functional genomics analysis shows that losses of complex morphological traits are associated with genome-wide divergence of the *cis*-regulatory landscape that is involved in the development of the trait. With regard to loss of limbs in snakes and degeneration of eyes in subterranean mammals, we consistently found that hundreds of CNEs diverged in the trait-loss species (i) are associated with genes that have key roles in development and function of this trait, (ii) significantly overlap regulatory elements that are specifically active in the respective tissues, and (iii) exhibit a clear loss signature of relevant transcription factor binding sites. Together, our results provide the first genome-wide evidence that widespread divergence of the *cis*-regulatory landscape is a hallmark of lost complex phenotypes.

While previous studies did not detect increased sequence divergence of many limb enhancers in snakes^18,22,23^, we report for the first time hundreds of limb regulatory elements that are substantially and specifically diverged in snakes. It remains an open question to what extent mutations in these regulatory elements directly contributed to the loss of limbs during evolution and to what extent sequence divergence happened as a consequence of neutral evolution after limb loss. Considering our findings that several snake-diverged limb regulatory elements are associated with genes that cause limb truncations and other malformations when perturbed, it is likely that mutations in some of these diverged regulatory elements, especially those associated with major limb patterning genes, contributed to gradual limb reductions in ancestral snakes that eventually led to complete loss of limbs. Similarly, it is plausible that mutations in some of the eye regulatory elements, where we and others^79^ detected sequence divergence in subterranean mammals, contributed to the gradual process of eye degeneration. In summary, our study presents the first comprehensive picture of non-coding changes that may have contributed to limb loss and eye degeneration, and provides many promising candidate loci to further investigate the molecular and developmental mechanisms underlying the loss of these traits.

Despite widespread divergence of limb regulatory elements in snakes, we also found many CNEs that overlap limb regulatory elements but are not substantially diverged in snakes, even after >100 My of evolution. One likely explanation for the maintenance of limb enhancers in limbless species is pleiotropy^23^. Notably, a previous study showed that a substantial portion of limb enhancers are also active in the developing genitals^22^, implying that selection for a genital-related function may preserve many limb regulatory regions. Indeed, comparing limb regulatory elements that are conserved or diverged in snakes, we found that conserved elements have a two-fold higher tendency to overlap pleiotropic regulatory elements that are also active in genitals (Supplementary Table 16), which supports that pleiotropy is a major factor for sequence conservation. Despite being largely conserved, such pleiotropic enhancers may nevertheless lose their limb regulatory activity. This was shown for the *Tbx4* HLEB enhancer and the Prox and Island I enhancers regulating *HoxD* genes, which lost enhancer function in the developing limbs but preserve activity in the developing genitals^22,88^. While our genome-wide CNE screen detected sequence divergence signatures of the Island I enhancer, the sequence changes that underlie such partial enhancer activity differences may often be subtle without significantly increasing overall sequence divergence (Supplementary Figure 10). This implies that the true extent of regulatory divergence in snakes is likely higher than what we report here.

Interestingly, our analysis not only detected divergence signatures related to limb loss and eye degeneration, but also signatures related to other phenotypes that changed in snakes and vision-impaired subterranean mammals. For example, we found that snake-diverged CNEs significantly overlap eye regulatory elements (Figure 2B; Supplementary Table 8). Together with the loss of opsins early in snake evolution^89^, this enrichment is consistent with a possible subterranean origin of the lineage. Furthermore, CNEs diverged in subterranean mammals are significantly associated with genes that lead to heart beat irregularities in a mouse knockout (“atrial fibrillation”, Supplementary Table 12). Indeed, irregularities in heart rhythmicity are known for the blind mole rat^90,91^ and a relative of the star nosed mole (European mole,^92^). Overall, these additional enrichments indicate signatures of regulatory divergence underlying other morphological and physiological changes in snakes and subterranean mammals.

Our study suggests a general strategy to detect sequence changes in ancestral regulatory elements that potentially contribute to morphological changes. By contrasting species with and without a particular phenotype, comparative genomics screens enable the systematic identification of non-coding genomic regions that exhibit specific divergence patterns and thus may exhibit differences in regulatory activity. Leveraging functional annotations and phenotypic data from gene knockouts in model organisms, enrichment tests can reveal associations between diverged genomic regions and specific functional categories or pathways, and thus highlight those regions that are associated with promising genes. Lastly, functional genomics can further prioritize these genomic regions based on overlap with regulatory elements that are active in relevant tissues. Approaches like ATAC-seq that require only small amounts of tissue make it now possible to also determine regulatory elements in nonmodel organisms, as shown here for tegu lizard embryos. Together, such a combined strategy has the potential to provide insights into the genomic basis of many morphological changes, which is fundamental to unravel the mechanisms underlying nature’s phenotypic diversity.

## Materials and Methods

### Tegu lizard and mouse tissue

The work with the tegu lizard was done in accordance with the Brazilian environmental and scientific legislation, under the SISBIO (Sistema de Autorização e Informação em Biodiversidade, Instituto Chico Mendes de Conservação da Biodiversidade) license 30309-4. Work with mouse was performed in accordance with the German animal welfare legislation and in strict pathogen-free conditions in the animal facility of the Max Planck Institute of Molecular Cell Biology and Genetics, Dresden, Germany. Protocols were approved by the Institutional Animal Welfare Officer (Tierschutzbeauftragter), and necessary licenses were obtained from the regional Ethical Commission for Animal Experimentation (Landesdirektion Sachsen, Dresden, Germany).

### Whole-genome sequencing and assembly of the tegu lizard

To sequence the genome of the tegu lizard *Salvator merianae*, we used tissue from an adult animal collected in Mato Grosso, Brazil, and deposited on the tissue bank collection of the Zoology Department from University of São Paulo (specimen accession number LG2117).

#### Whole-genome sequencing

Genomic DNA was extracted from liver tissue of an adult animal following a standard phenol-chloroform extraction protocol. To generate fragment libraries, purified DNA was subjected to standard Illumina DNA library preparation. In brief, DNA was sheared for 550 bp by sonication (Covaris S2). After XP bead purification (Beckman Coulter), ends were polished and A-tailed and universal adapters were ligated (TruSeq DNA PCR-Free kit, Illumina). After ligation, adapters were depleted by XP bead purification (Beckman Coulter). Sample indexing was done during PCR (15 cycles). Libraries were quantified with Fragment Analyzer (Advanced Analytical) and KAPA (KAPA Biosystems), and equimolar pooled for sequencing. To generate mate-pair libraries, purified DNA was subjected to the Nextera Mate Pair Library Preparation protocol (Illumina). For accurate sizing after tagmentation, gel-based size selections for an average of 2 Kb and 10 Kb were done. The resulting libraries were quantified with Fragment Analyzer (Advanced Analytical) and KAPA (KAPA Biosystems) and equimolar pooled for sequencing. We sequenced 2x300 bp reads from three fragment libraries on the Illumina MiSeq platform, and 2x150 bp and 2x100 bp reads from two 2 Kb and two 10 Kb mate-pair libraries on the Illumina HiSeq 2500 platform to a total sequencing coverage of 104X (Supplementary Table 1).

#### Pre-processing of sequencing reads

Raw reads were trimmed for the presence of sequencing adapters by applying cutadapt^93^, setting a minimum read length of 100 bp for fragment reads and 50 bp for mate-pair reads. Trimmed reads were further processed by permuting ambiguous bases using SGA^94^ (parameters ‘*sga preprocess --permute ambiguous*’). The resulting reads were corrected for sequencing errors applying the SGA-ICE correction pipeline^25^, running iterative k-mer-based corrections with increasing k-mer sizes (k=40, 100 and 200 for MiSeq reads; k=40, 70, 100, 125, 150 for HiSeq reads).

#### Genome assembly

Error-corrected reads were assembled using ALLPATHS-LG^26^. We set CLOSE_UNIPATH_GAPS=False to circumvent exceedingly long runtimes related to the relatively long 300 bp fragment reads, and HAPLOIDIFY=True to account for the fact that we are assembling reads from diploid organisms. ALLPATHS-LG performs all assembly steps, including contig and scaffold assembly. The resulting 2.026 Gb assembly comprises 5,988 scaffolds, with 4% of the bases located in assembly gaps.

#### Assessing assembly completeness by mapping Ultra Conserved Elements (UCEs)

We obtained a set of non-exonic ultraconserved elements (UCEs) that are deeply conserved across vertebrates and should thus be found in a correctly assembled genome. Specifically, from all 481 UCEs (originally defined as genomic regions of at least 200 bp that are identical between human, mouse and rat^27^), we first excluded all UCEs that overlap exons in human (UCSC “ensGene” gene table^95^). Then, we used liftOver^95^ (parameters ‘*-minMatch=0.1*’) to obtain those UCEs that align to chicken (galGal5 assembly), zebrafish (danRer10 assembly) and medaka (oryLat2 assembly), resulting in 197 vertebrate non-exonic ultraconserved elements. We used lastz^96^ (parameters ‘*--gappedthresh=3000 --hspthresh=2500 --seed=match6*’) to align these sequences to our tegu genome assembly and to other reptiles. All alignments were filtered for a minimum of 75% identity and a length of at least 50 bp. The number of aligning UCEs is reported in Supplementary Table 2.

### Tegu transcriptome sequencing and assembly

#### Transcriptome sequencing

We extracted total RNA from two tegu lizard embryos around stage DH28-30, which corresponds to mouse embryonic stage E11-E11.5 ^97^. Tissues were immediately frozen in liquid nitrogen and total RNA was later extracted following a standard Trizol extraction. For the generation of mRNA libraries, mRNA was isolated with the NEBNext Poly(A) mRNA Magnetic Isolation Module according to the manufacturer’s instructions. After chemical fragmentation, the samples were directly subjected to strand specific RNA-Seq library preparation (Ultra Directional RNA Library Prep, NEB). For adapter ligation, custom adaptors were used (Adaptor-Oligo 1: 5′-ACA-CTC-TTT-CCC-TAC-ACG-ACG-CTC-TTC-CGA-TCT-3′, Adaptor-Oligo 2: 5′-P-GAT-CGG-AAG-AGC-ACA-CGT-CTG-AAC-TCC-AGT-CAC-3′). After ligation, adapters were depleted by XP bead purification (Beckman Coulter). Sample indexing was done during PCR enrichment (15 cycles). The resulting libraries were quantified with Fragment Analyzer (Advanced Analytical) and equimolar pooled for sequencing. We sequenced 2x75 bp reads from eight strand-specific mRNA libraries on the Illumina HiSeq 2500 platform (Supplementary Table 1).

#### Pre-processing of sequencing reads and transcriptome assembly

We trimmed raw reads for the presence of sequencing adapters with cutadapt^93^, setting a minimum read length of 30 bp. Trimmed reads were mapped against the tegu genome using TopHat2^98^ (parameters ‘*--library-type fr-firststrand*’) and assembled using Cufflinks^99^ (parameters ‘*--library-type fr-fìrststrand’*). In addition, we also performed a *de novo* transcriptome assembly with Trinity^100^ (parameters -- *SS_lib_type FR --genome_guided_max_intron 10000*’).

### Sequencing of open chromatin for tegu lizard and mouse

#### ATAC sequencing

ATAC-seq^35^ was performed on 10-day old tegu lizard embryos (DH29-30; corresponding to mouse E11.5), and on E11.5 and E14.5 mouse embryos. Tegu lizard embryos were dissected in PBS to obtain samples from limb, brain, heart, liver, tissue flanking the limb buds and the remaining embryonic material. Mouse embryos were dissected to obtain whole eye, limb and midbrain at E11.5, and lens and retina separately at E14.5. For both tegu lizard and mouse embryos, tissue samples were pooled from at least 10 embryos and frozen in liquid nitrogen immediately after dissection. ATAC was performed on each tissue in at least two biological replicates (Supplementary Table 1). For each sample, we ground approximately 4 mg of tissue on a glass douncer using a milder lysis buffer containing 10% of the recommended detergent concentration. Amplification of ATAC libraries was done for 10 PCR cycles with Illumina Nextera reagents. The resulting libraries were quantified with Fragment Analyzer (Advanced Analytical) and KAPA (KAPA Biosystems) and equimolar pooled for sequencing. We sequenced 2x75 bp reads on the Illumina HiSeq 2500 and NextSeq platform (Supplementary Table 1).

#### Pre-processing of sequencing reads

We trimmed raw reads for the presence of sequencing adapters with cutadapt^93^, setting a minimum read length of 30 bp. Trimmed reads were mapped against our tegu assembly or the mouse mm10 genome using Bowtie2^101^ (default parameters). To center reads at the actual transposase insertion site, we shifted reads that mapped to the plus strand by +4 bp, and reads that mapped to the minus strand by −5 bp, as described in^35^.

#### Peak calling and identification of differential peaks

We used MACS2^102^ to identify discrete peaks of enriched ATAC-seq signal, using the mappable portion of the tegu genome (Hotspot getMappableSpace.pl script;^103^) and using the shifted reads as input (MACS2 parameters ‘*--nomodel --shift −50 --extsize* 100’). Finally, peaks that show differential signal across tissues were identified using DiffBind^104^ (parameters ‘*method=DBA_EDGER, bFullLibrarySize=FALSE, bSubControl=FALSE, bTagwise=FALSE*’; fold change equal or greater than 1).

### Tegu genome annotation

#### Repeat masking

We first applied RepeatModeler (http://www.repeatmasker.org/; parameters ‘*-engine ncbi*’) to *de-novo* identify repeat families in the tegu genome. Then, we used RepeatMasker (default parameters) with the resulting repeat library to soft-mask the tegu genome.

#### Gene annotation

In order to annotate the tegu genome, we prepared the following four datasets:

1. Assembled tegu transcriptome: Cufflinks genome-guided transcriptome assembly (72,538 transcripts; see above) and Trinity *de novo* transcriptome assembly (6,156 transcripts; see above).
2. Mapping proteins of closely-related species to the tegu genome: We mapped Ensembl/UniProt protein sequences from the green anole lizard *Anolis carolinensis*, and the turtles *Pelodiscus sinensis* and *Chelonia mydas* to the soft-masked tegu genome using Exonerate^105^ (parameters ‘*-m protein2genome --exhaustive 0 --subopt 0 --bestn 1 --minintron 20 --maxintron 50000 --softmasktarget T --refine region -- proteinhspdropoff 20*’).
3. Augustus gene models: To use Augustus^106^ for gene prediction, we first constructed a training set of gene models from the cufflinks-assembled transcripts and Trinity transcript contigs using the PASA pipeline^107^. Trinity contigs were preprocessed with SeqClean (https://sourceforge.net/projects/seqclean/) to remove polyA tails, vector/adapter sequences and low-complexity regions. Next, with the PASA pipeline, we mapped the contigs to the soft-masked genome with GMAP^108^ and BLAT^109^ and removed low-quality alignments (alignment identity less than 90%, or less than 70% aligned). We then combined the remaining Trinity contig alignments with the cufflinks-assembled transcripts by collapsing redundant transcript structures and clustering overlapping transcripts. Finally, we used TransDecoder (https://github.com/TransDecoder/TransDecoder) to search within the resulting PASA transcripts for candidate coding regions, keeping only those transcripts, which encode a complete open reading frame (with start and stop codon) of at least 100 aa. Using this set of gene models as training data, we trained Augustus on the tegu genome after hard-masking interspersed repeats.
4. CESAR gene mappings: We used CESAR^110,111^, a Hidden Markov Model based exon aligner that takes reading frame and splice site information into account, and our genome alignment between tegu and human (see below) to map human protein-coding genes to the tegu genome. Using 196,269 exons from 19,919 human protein-coding genes (Ensembl78, longest isoform per gene), CESAR annotated 162,959 exons with an intact reading frame and consensus splice sites belonging to a total of 17,053 genes in the tegu lizard.

Using these four datasets, we annotated genes in the tegu genome with MAKER^112^. Cufflinks-assembled transcripts were passed onto MAKER via the *est_gff* option on maker_opts.ctl file. The Ensembl/UniProt proteins from anole lizard and two turtles were passed via the *protein_gff* option. Augustus gene predictions were combined with CESAR mappings and were passed to MAKER via the *pred_gff* option. Furthermore, we specified the Augustus gene models in *augustus_species* in maker_opts.ctl file. The options *est2genome, protein2genome*, and *keep_preds* in maker_opts file were all set to 1. We ran a single iteration of MAKER, resulting in 70,462 genes and 70,486 transcripts. After filtering out all entries with an Annotation Edit Distance (AED) value of 1, the final gene annotation for the tegu genome contains 23,463 genes and 23,487 transcripts.

To obtain tegu-human orthologs, we made use of the CESAR mapping of human genes to the tegu genome as follows. First, we removed single-exon mappings shorter than 100 bp and mappings which result in very large introns (longer than 200 Kb), provided that the length of the corresponding intron in the human Ensembl transcript is greatly different (200 Kb difference). We also filtered out those MAKER gene predictions which correspond to <100 bp single-exon genes. Next, we intersected the cleaned CESAR and MAKER gene sets to obtain those tegu genes which correspond to a human gene. This resulted in a set of 16,284 tegu genes with a human ortholog.

### Multiple genome alignments

Genome alignments were done as described in^113,114^. To analyze limb loss in snakes, we used lastz^96^ (parameters ‘*K=2400 L=3000 Y=3000 H=2000’* and the HoxD55 scoring matrix) to compute pairwise local genome alignments between tegu and 28 other vertebrate genome assemblies (Supplementary Table 3). We used axtChain^115^ (default parameters) to build co-linear alignment chains. For species other than the more closely-related squamates (green anole and dragon lizard, boa and python), we subsequently used more sensitive local alignments to find additional co-linear alignments by running lastz (parameters *K=2000 L=2700 W=5’*) on all chain gaps (non-aligning regions flanked by local alignments) that are between 20 bp and 100 Kb long. All local alignments were quality-filtered by keeping only those alignments where at least one ≥30 bp region has ≥60% sequence identity and ≥1.8 bits entropy as described in^113^. We used chainNet^115^ (default parameters) to obtain nets from a set of chains. To remove low-scoring alignment nets that are unlikely to represent real homologies, we filtered these nets for a span in the reference and query genome of at least 4 Kb and a score of at least 20,000 (for frog and coelacanth we required a score of at least 10,000). In addition, we kept all nets that represent inversions or local translocations if they score at least 10,000. The filtered pairwise alignment nets are the input to Multiz^116^ to build a multiple alignment. The phylogenetic position of squamate species was taken from^117^.

To analyze loss of vision in mammals, we used the same lastz/chain/net/Multiz pipeline (parameters as above) to align 20 mammals, green anole lizard, chicken and frog (Supplementary Table 9) to the mouse mm10 assembly, which serves as the reference. We kept alignment nets with a minimum score of at least 70,000 and a minimum span of 9 Kb, or a minimum score of 150,000 and a minimum span of 6 Kb.

For the three non-mammalian species, we used the additional round of sensitive local alignments, as described above, and kept alignment nets with a score of at least 15,000.

### Detection of Conserved Non-coding Elements (CNEs) diverged in species of interest

#### CNE identification

To detect evolutionarily conserved elements in the tegu and mouse multiple alignment, we used PhastCons^118^ (parameters *‘expected-length=45, target-coverage=0.3 rho=0.3’*). Since PhastCons also requires a phylogenetic tree with neutral branch lengths as input, we estimated branch lengths from 4-fold degenerated third codon positions using phyloFit^118^ (parameters *‘--EM--precision HIGH --subst-mod REV)*. We combined the results of PhastCons with conserved regions detected by GERP^119^ (default parameters). In addition, we required that conserved regions align well to at least 4 of the 14 mammals in the tegu-based alignment, and at least 15 of all species in the mouse-based alignment.

To obtain CNEs from the full set of conserved elements, we stringently excluded all conserved parts that overlap potentially coding regions. For the tegu genome, we obtained a comprehensive set of potentially coding sequences from four sources. First, we used proteins from the green anole lizard and the turtles *Pelodiscus sinensis* and *Chelonia mydas* that were mapped with Exonerate (see above). Second, we downloaded Refseq proteins (“vertebrate_other”) from NCBI and mapped them to the tegu genome with blastx^120^ (E-value cutoff 0.01). Third and fourth, we added the full sets of CESAR human mappings and MAKER gene predictions (see above). For the well-annotated mouse genome, we obtained all coding exons listed in the UCSC tables “ensGene”, “refGene” and “knownGene”^95^. A 50 bp flank was added to both sides of all coding sequences to also exclude conserved splice site regions. To obtain CNEs, we subtracted the respective coding bases from all conserved elements. Finally, we selected those CNEs that are at least 30 bp long.

#### Measuring sequence divergence of CNEs

To compute the percent identity between the CNE sequence of a species and the reconstructed ancestral sequence, we used the previously developed pipeline described in^80^. This pipeline uses the phylogeny-aware PRANK alignment method^121^ (parameters *‘-keep -showtree -showanc -prunetree -seed=10’*) to align the sequences of the extant species and to reconstruct the most likely ancestral sequences of each CNE. It then computes the percent sequence identity between the aligned ancestral sequence and the CNE sequence of each species^80,122^.

#### Detection of CNEs diverged in snakes

To account for lineage-specific evolutionary rates, the resulting percent sequence identities were normalized by the total branch length between each species and the reconstructed ancestor:

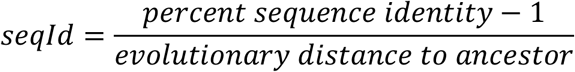

Sequence identity values are thus in the negative range: the lower the sequence identity, the higher the sequence divergence, and vice versa. Both boa and python snakes belong to the same lineage, representing a single evolutionary loss. We therefore calculated a Z-score for each CNE, comparing the sequence identity value of the least diverged snake with the average and standard deviation of sequence identity values of all limbed species:

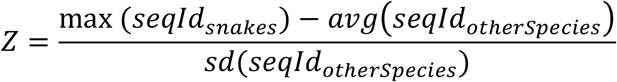

To obtain CNEs specifically diverged in snakes, we first considered only those CNEs for which sequence identity values were calculated for both snakes, all three lizards, gecko, and at least 3 additional amniote species. This excludes CNEs that have missing genomic data for too many species. Second, we calculated a global Z-score by comparing snakes with all other species, and removed those CNEs with a global Z-score ≥ −3. Finally, we calculated a local Z-score by comparing snakes with tegu, green anole and dragon lizards, and removed CNEs with a local Z-score ≥ −3. This resulted in 5,439 snake-diverged CNEs used for further analysis.

We also generated a control set of diverged CNEs using the same Z-score calculation but considering the green anole lizard and dragon lizard as trait-loss species. Using a Z-score cutoff of −3, this resulted in 614 CNEs that are more diverged in the two limbed lizards compared to all other species. To rule out that a lack of significant enrichments is due to the lower number of CNEs in this control set, we also obtained and tested a second control set consisting of the same number of CNEs (the top 5,439 ranked by the global Z-score and requiring a local Z-score < −3).

#### Detection of CNEs diverged in subterranean mammals

In contrast to snakes, the degeneration of the visual system has happened independently in four subterranean mammals. Therefore, we applied the branch method of the “Forward Genomics” approach^80^ to select CNEs that are significantly more diverged in the four vision impaired species compared to all other species. The branch method considers both phylogenetic relatedness between species and differences in their evolutionary rates, and computes the significance of the association between the sequence divergence and phenotype. CNEs for which sequence identity values could not be calculated for all four vision impaired species were excluded. The resulting p-values were corrected for multiple testing using the Benjamini & Hochberg method^123^. Using an adjusted *p*-value cutoff of 0.005, this resulted in 9,364 diverged CNEs used for further analysis.

We generated a control set of diverged CNEs using the same Forward Genomics setup but considering human, rat, cow, and elephant as trait-loss species. Using the same adjusted *p*-value cutoff, this resulted in 86 CNEs that are more diverged in these four mammals compared to all other species. As for snake-diverged CNEs, we also obtained a second control set with the same number of CNEs by taking the top-ranked 9,364 CNEs.

### Functional enrichment analysis

We statistically tested if diverged CNEs are significantly associated with genes belonging to a functional class, using all remaining non-diverged CNEs as a control. To associate CNEs to potential target genes, we used the regulatory domain concept of GREAT^32^. Briefly, for each gene, we defined a basal (promoter-associated) domain of 5 Kb upstream and 1 Kb downstream of the transcription start site and a distal domain that extends the basal domain up to the basal domain of the next gene or at most 300 Kb in either direction (Supplementary Figure 3). Transcription start sites for the mouse mm10 genome were downloaded from the GREAT website. The overlap of a CNE with one or more regulatory domains is then used to associate the CNE to its target gene(s).

To obtain functional annotations for all genes genome-wide, we used the Mouse Genome Informatics (MGI) Phenotype ontology^34,124^ that lists phenotypes observed in mouse gene knockouts (MGI table MGI_PhenoGenoMP.rpt). We only considered phenotypes resulting from single gene knockouts. All phenotypes associated to at least 3 genes were tested for enrichment. For the snake-diverged CNE analysis, we mapped MGI phenotypes to the corresponding tegu genes by using our CESAR annotation (see above). We first tested enrichments of snake-diverged CNEs using all mouse knockout phenotypes. However, this analysis was not meaningful as the top enriched terms were biased towards mammalian-specific characteristics such as placenta, milk, brown fat, or testicular descent, which reptiles never possessed. Therefore, we specifically tested enrichment of limb MGI knockout phenotypes (all child terms of MP:0002109 “abnormal limb morphology”). In addition to MGI annotations, we used a set of 435 genes that are involved in limb patterning, growth and tissue differentiation^33^. For the analysis of CNEs diverged in vision-impaired subterranean mammals, we tested all MGI knockout phenotypes.

For each gene set (genes with the same functional annotation), we used the LOLA R package ^125^, which implements a one-sided Fisher’s exact test, to ask whether diverged CNEs overlap the regulatory domains of this gene set more often than the remaining non-diverged CNEs. To exclude the possibility that significant results are explained by several CNEs that are in close proximity, we merged all CNEs closer than 50 bp into one CNE for these tests, reducing the number of diverged CNEs to 4,849 for the limb study and 9,259 for the eye study. The resulting p-values were corrected for multiple testing using the Benjamini & Hochberg method^123^. To further estimate the magnitude of an enrichment as a Z-score, we randomly subsampled, 10,000 times, same-sized sets from the full set of CNEs and determined the overlap with regulatory domains of genes in this set.

To test if diverged CNEs significantly overlap regulatory elements active in a particular tissue, we used LOLA and random subsampling as we did for regulatory domains of gene sets (see above). In addition to our ATAC-seq peaks, we obtained a large number of publicly available regulatory datasets^22,33,40,48–52,86,87,126–131^ to get tissue-specific peaks. In short, for each experimental assay, we selected peaks observed in at least two tissue/stage replicates (when available), and obtained tissue-specific peaks by subtracting peaks observed with this assay in other tissues (when available). To test for enrichments of snake-diverged CNEs, coordinates of regulatory elements were mapped to the tegu genome^95^ (liftOver, parameters *‘-minMatch=0.1’*). Details about all limb and eye datasets, the underlying experiments, tissues, time-points, and species, and the overlap statistics used for LOLA’s one-sided Fisher’s exact test are provided in Supplementary Tables 8 and 15.

### Scoring transcription factor binding sites

A motif library was generated by integrating motifs from TRANSFAC^132^, JASPAR^133^ and UniPROBE^134^. We clustered the resulting 2,272 motifs into 638 groups to reduce motif redundancy based on a similarity score computed with Tomtom (parameters ‘*-thresh=1-dist=ed*) (Supplementary Table 17). We then scored each CNE sequence of each species for the presence of each of the motifs by applying SWAN^135^ (parameters *‘window length 200 bp; GC-matched random generated background sequence; -wc’*), a Hidden Markov Model based method that considers both weak and strong motif occurrences and avoids inflated scores in case motif occurrences overlap each other. Since SWAN scores do not consider the exact position of each motif and since we consider the sequence of each species separately, our results are not affected by turnover of the same motif in the CNE sequence.

To determine if sequence divergence resulted in the loss of well-conserved TF motifs, we first selected those CNE-motif pairs, for which the reconstructed ancestral sequence had a score exceeding random expectation. Since different transcription factors have different binding strengths, we calculated a stringent TF-specific score cut-off that indicates the presence of at least one TF motif in an otherwise random sequence. To this end, we generated 100 random sequences of length 200 bp each for 21 GC-content bins that range from 0 to 100%. Then, we inserted a binding site (sampled from the motif) into the random sequences, applied SWAN, and determined the 90% score quantile of each bin. This 90% score quantile corresponds to a GC-dependent motif cut-off where 10% of the GC-matched random sequences with the inserted motif have a score at least as high. Then, we only considered CNE-motif pairs, for which the reconstructed ancestral sequence scored above this threshold. For those CNEs, we then determined the score of this motif for the CNE sequence of each species.

For the limb study, we compiled a list of 65 transcription factors that are known to function in limb development (Supplementary Table 18). Our motif library contains 55 motifs for these TFs (Supplementary Table 18). CNEs were separated into two groups: (i) snake-diverged CNEs that overlap limb-specific regulatory data (Supplementary Table 4), and (ii) all other CNEs. A one-sided Wilcoxon rank sum test was used to assess if diverged CNEs have significantly higher motif scores than the other CNEs. Resulting p-values were corrected for multiple testing using the Benjamini & Hochberg method^123^. We also computed the median score for CNE-motif pairs separately for both CNE groups and visualized the difference in the median in Figure 4A. Accordingly, for the eye study, we compiled a list of 60 transcription factors that are known to function in eye development, resulting in 58 motifs contained in our library (Supplementary Table 19) and visualized the difference in the median between CNEs diverged in subterranean mammals that overlap eye-specific regulatory data (Supplementary Table 10) and all other CNEs in Figure 4C.

## Data availability

All primary data, including the tegu lizard genome assembly, its gene annotation, CNEs, ATAC-seq peaks, the tegu multiple genome alignment and the mouse multiple genome alignment are available at XXXXXXXX. All Illumina sequencing data will be uploaded to NCBI.

## Code availability

Our study used publicly available software. All tools and methods, their version number, and the used parameters are detailed in Supplementary Table 20.

## Acknowledgments

We thank the genomics community for sequencing and assembling the genomes of the many vertebrates used here, and the UCSC genome browser group for providing software and genome annotations. We also thank Virag Sharma for running CESAR, Nadine Vastenhouw for experimental infrastructure, Terence Capellini for helpful comments on the manuscript, Jared Simpson for help with SGA, Andreas Dahl and Sylke Winkler for DNA sequencing, the Computer Service Facilities of the MPI-CBG and MPI-PKS, and the DNA Sequencing, Microarray and Biomedical service facilities of the MPI-CBG for their support. This work was supported by the Max Planck Society, and by FAPESP stipends 2012/01319-8 and 2012/23360 to JGR.

